# Juvenile hormone affects the development and strength of circadian rhythms in young bumble bee (*Bombus terrestris*) workers

**DOI:** 10.1101/2020.05.24.101915

**Authors:** Atul Pandey, Uzi Motro, Guy Bloch

**Affiliations:** Department of Ecology, Evolution, and Behavior, The Hebrew University of Jerusalem, Jerusalem, Israel; The Federmann Center for the Study of Rationality, The Hebrew University of Jerusalem, Jerusalem, Israel

**Keywords:** Hormone, Juvenile hormone, bumble bee, *Bombus terrestris*, circadian rhythms, locomotor activity, sleep, gonadotropin

## Abstract

The circadian and endocrine systems influence many physiological processes in animals, but little is known on the ways they interact in insects. We tested the hypothesis that juvenile hormone (JH) influences circadian rhythms in the social bumble bee *Bombus terrestris*. JH is the major gonadotropin in this species coordinating processes such as vitellogenesis, oogenesis, wax production, and behaviors associated with reproduction. It is unknown however, whether it also influences circadian processes. We topically treated newly-emerged bees with the allatoxin Precocene-I (P-I) to reduce circulating JH titers and applied the natural JH (JH-III) for replacement therapy. We repeated this experiment in three trials, each with bees from different source colonies. Measurements of ovarian activity confirmed that our JH manipulations were effective; bees treated with P-I had inactive ovaries, and this effect was fully reverted by subsequent JH treatment. We found that JH augments the strength of circadian rhythms and the pace of rhythm development in individually isolated newly emerged worker bees. JH manipulation did not affect the free-running circadian period, overall level of locomotor activity, or the amount of sleep. Given that acute manipulation at an early age produced relatively long-lasting effects, we propose that JH effect on circadian rhythms is mostly organizational, accelerating the development or integration of the circadian system.

## Introduction

The circadian and endocrine systems are pivotal for the integration of external and internal information and for coordinating processes in multiple tissues (Bedrosian et al., 2016; Neumann et al., 2019; Tsang et al., 2013). In vertebrates, particularly in mammals, there is good evidence that these two regulatory systems interact. The circadian system influences endocrine tissues and processes, resulting in circadian rhythms in the circulating levels of many vertebrate hormones (Bedrosian et al., 2016; Hastings et al., 2007; Kriegsfeld et al., 2002; Neumann et al., 2019; Tsang et al., 2013). Hormones also influence circadian functions. For example, gonadal hormones may lengthen the free-running period, decrease period precision, or reduce the duration of daily activity bouts (e.g., Albers, 1981; Daan et al., 1975; Iwahana et al., 2008; Jechura et al., 2000; Karatsoreos et al., 2011, 2007). Many of these effects are at least partially mediated by hormone receptors in the suprachiasmatic nucleus (SCN), the central brain clock of mammals (He et al., 2007; Iwahana et al., 2008; Karatsoreos et al., 2011, 2007; Sellix et al., 2004). Steroid hormones, including progestins, corticosteroids, estrogens, and androgens, were also shown to influence circadian rhythms in locomotor activity and transcription levels of core circadian clock genes in fishes (Zhao et al., 2018).

The interplay between the circadian and endocrine systems is relatively little explored in adult insects (Bloch et al., 2013). Only a few studies recorded hormone titers throughout the day under constant conditions. Nevertheless, these measurements, together with indirect evidence for circadian modulation of hormone biosynthesis rate, and the expression of genes encoding proteins involved in hormone biosynthesis, hormone binding, or hormone degradation, suggest that the circadian system influences the circulating levels of many insect hormones. There is also little evidence for hormonal regulation of circadian rhythms in insects (reviewed in Bloch et al., 2013). This includes the best-studied insect hormone, juvenile hormone (JH), which functions as a gonadotropin in many insects. There is some evidence that JH and ovarian activity influences circadian rhythms in the cockroach *Blattella germanica*, although the data is quite perplexing. Females of this species show strong circadian rhythms during the vitellogenic phase of the reproductive cycle when JH titers are expected to be high, but not in sexually receptive females during the first gonadotropic cycle (Lee and Wu, 1994). Active ovaries mask the expression of circadian rhythms, and allatectomy abolished the strong circadian rhythms that are shown by ovariectomized females. However, replacement therapy with a JH analog did not restore circadian rhythmicity (Lin and Lee, 1998).

Mutations in the JH receptors Met, and the JH biosynthesis pathway enzyme JH acid O-methyltransferase (JHAMT) attenuated the strength of circadian rhythms in *D. melanogaster*, suggesting that JH affects circadian rhythms in this species too (Wu et al., 2018). Studies with Milkweed bug, *Oncopeltus fasciatus* in which JH levels were decreased by treatment with Precocene-II, and increased by JH supplementation produced conflicting evidence concerning the effects of JH on the circadian rhythm of feeding and mating behavior (Walker, 1977; Woodard and Rankin, 1980). JH does not seems to be involved in the regulation of circadian rhythms in the cockroach *Blaberus discoidalis* because circadian rhythms in locomotor activity were similar in the allatectomized and control-treated individuals (Shepard and Keeley, 1972). The influence of JH on circadian rhythms was also studied in the Western honey bee *Apis mellifera* in which JH does not function as a major gonadotropin. JH manipulation by allatectomy and replacement therapy with the JH analog methoprene, which successfully affected the age of first foraging (Sullivan et al., 2000), failed to affect circadian rhythms in locomotor activity (Bloch et al., 2002). Similar JH manipulations did not have a consistent influence on the circadian brain expression of the canonical clock gene *Period*. However, there was a trend towards aberrant cycling in allatectomized bees (Bloch and Meshi, 2007).

Taken together, the available studies revealed significant variability in the effects of JH manipulation on circadian rhythms in insects and suggest that some of this variability may relate to whether or not JH functions as a gonadotropin. To test this hypothesis, we studied the influence of JH on circadian rhythms in the social bumble bee *Bombus terrestris*. Bumble bees are taxonomically related to honey bees, but live in smaller annual colonies, showing a simpler form of social organization (Michener, 1974). By contrast to honey bees, in bumble bees, JH is the major gonadotropin (Shpigler et al., 2014), and does not affect task performance (i.e., brood care vs. foraging activity; (Shpigler et al., 2016). However, in both species, the division of labor is similarly correlated with the expression of circadian activity rhythms; foragers have strong circadian rhythms, whereas nurse bees are typically active around the clock with attenuated rhythms (reviewed in Bloch, 2010; Eban-Rothschild and Bloch, 2012). We manipulated JH levels in young bumble bee workers using topical treatments with the allatoxin precocene 1 (P-I) to reduce hemolymph JH titers and the natural JH of bumble bees (JH-III; Bloch et al., 2000, 1996) for replacement therapy. Our results suggest that JH affects the strength and development of circadian rhythm in locomotor activity in *Bombus terrestris*.

## Materials and Methods

### Bees

Colonies of *Bombus terrestris* containing a queen, 5-10 workers, and brood at various developmental stages (typically 2–4 days post first worker emergence) were purchased from Polyam Pollination Services, Kibbutz Yad-Mordechai, Israel (Trial 1 & 2) or BioBee Biological systems Ltd. Kibbutz Sde Eliyahu, Israel (Trial 3). We housed each colony in a wooden nesting box inside an environmental chamber (29 ± 1 °C; 55% ± 10% RH) in constant darkness at the Bee Research Facility at the Edmond J. Safra Campus of the Hebrew University of Jerusalem, Givat Ram, Jerusalem. The nest boxes (21 × 21 × 12 cm) were made of wood and were outfitted with a top and front wall made of transparent Plexiglas panels. The colonies were fed *ad libitum* with commercial sugar syrup and pollen cakes made of honey bee collected pollen that was purchased from Polyam Pollination Services. We performed all treatments and feeding under dim red light (DD; using Edison Federal EFEE 1AE1 Deep Red LED; mean wavelengths = 740 nm, maximum and minimum wavelengths = 750 and 730, respectively) minimizing the production of substrate-born vibrations.

### General experimental outline

Our experimental design is summarized in **Fig. 1**. At day-1 of each trial, we collected 120–130 newly-emerged worker bees (<24h of age) from a pool of 12-15 ‘donor colonies’, chilled them on ice, measured their thorax width, treated them as detailed below (~30 bees per treatment under dim red light), and placed them in groups of four of the same treatment in small wooden cages (12 × 5 × 8 cm). The bees were collected in two batches such that a bee was chilled on ice for no longer than 90 min, and the entire procedure took 2–3 hours from collection in the colony until the end of the first treatment (precocene-I, castor oil or control handling, see **Fig. 1** and below). On Day 3, approximately 24–26 h after the second treatment, we transferred the bees to an environmental chamber, isolated them each in an individual cage, and monitored their locomotor activity for 5–9 consecutive days (five days in Trial 1, eight days in Trial 2, and nine days in Trial 3, see **Fig. 1**) under tightly controlled environmental conditions (29 ± 1 °C; 55% ± 5% RH). At the end of the monitoring session, we stored all the live bees in a −20 °C until assessing their ovarian state. We repeated this experiment three times: in April 2017 (Trial 1), May 2017 (Trial 2), and April–May 2019 (Trial 3).

**Figure 1.**
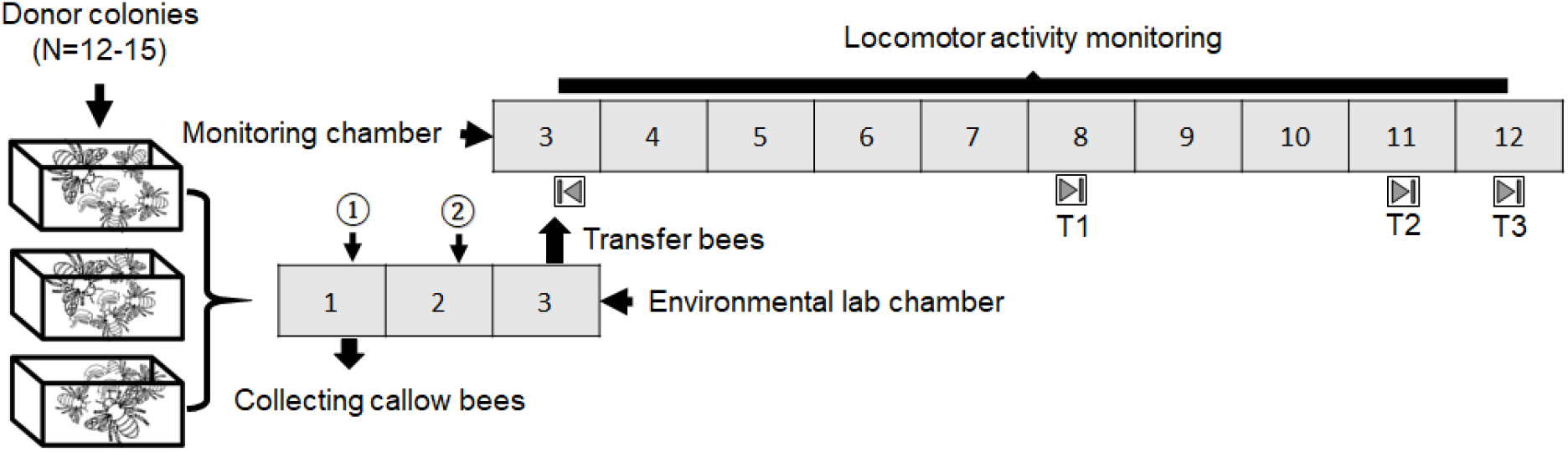
Schematic description of the general experimental outline. The numbers represent the day of the experiment measured as time passed after collecting the callow bees. On Day 1 between 14.00-15.00 hrs, we collected newly emerged (0–24hr of age) worker bees from 12–15 queenright “donor” colonies and subjected them to the first treatment at around 16:00. About 24 hrs later, on the afternoon of Day 2, we subjected the bees to the second treatment. In the afternoon (16:00-18:00) of Day-3, we transferred each bee into an individual monitoring cage and monitored locomotor activity for 5–9 successive days under contestant laboratory conditions. At the end of the monitoring session, we collected the bees and recorded their ovarian state. N=number, ①-first treatment, ②-second treatment, 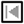-start, and 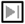-end of monitoring session trial; T1=trial-1, T2=trial-2, and T3-trial-3. All treatments, observations, and manipulations were conducted under dim red light.

### Manipulating circulating JH titers

We subjected the callow bees to one of four treatment groups. Control bees (*“Control”*), were handled and chilled on ice (20–25 min) similarly to bees from the other experimental groups, on both Day 1 and Day 2, but not treated with any drug or vehicle. Sham-treated bees (“*Sham*”), were chilled on ice and topically treated with castor oil on Day1, and with dimethylformamide (DMF, J.T Backers, cat # 7032; 3.5ul/bee, irrespective of body sizes) on Day 2. The amount of castor oil (Sigma-Aldrich, cat # 259853; 4.0-5.2μl/bee) was adjusted according to body size, as detailed in **Table 1**. To reduce JH levels, we treated the bees on Day 1 with the allatoxin Precocene-I (“*P-I*”; Sigma-Aldrich, cat # 195855) suspended in castor oil vehicle solution. We thoroughly mixed the P-I, and castor oils by repeated pipetting followed by vortexing the mixture at high speed for 2–3 min. We adjusted the P-I treatment to body size (200-260μg /4.0–5.2μl castor oil/bee; **Table 1**), as previously reported (Pandey et al., 2020). Replacement therapy (“*P-I+JH*”) treated bees were topically treated with P-I on Day 1, as described above. On the following day, we topically treated the bees with 50μg JH-III (Sigma-Aldrich, cat # J2000) dissolved in 3.5μl DMF vehicle solution. We handled and treated the control and sham control bees similarly to their respective treatment groups. Following the P-I, and JH-III treatments, we kept the bees on ice-chilled glass plates immobilized for ~10 minutes in order to improve drug absorption and minimize possible wiping-off. Additional details on our JH manipulation procedures, and the validation of our assays, can be found in Shpigler et al., (2016) and Pandey et al., (2020).

**Table 1:** Body size adjusted mounts of Precocene-I used to reduce JH titers in callow bees.

| Size | Head Width (mm) | Precocene-I (µg/bee) | Castor oil (µl/bee) |
| --- | --- | --- | --- |
| Small | 2.4-2.6 | 200 | 4.0 |
| Small-medium | 2.7-2.9 | 220 | 4.4 |
| Large-medium | 3.0-3.5 | 240 | 4.8 |
| Large | 3.5-4.2 | 260 | 5.2 |

### Monitoring locomotor activity

We placed each bee individually in a transparent monitoring cage (made of a modified 90 mm Petri dish). The cages with the focal bees were transferred to the monitoring environmental chamber inside a lightproof box to avoid any light exposure. The monitoring chamber was illuminated with a dim red light that the bees cannot see (Edison Federal EFEF 1AE1 Far [Cherry] Red LED; mean wavelength = 740 nm, maximum and minimum wavelengths were 750 and 730, respectively). The location of each focal bee was recorded with one of four CCD cameras (Panasonic WV-BP334, 0.08 lux CCD video cameras) and an image acquisition board (IMAQ 1409, National Instruments, Austin, TX) at a frequency of 1 Hz over the entire monitoring session as previously described (Shemesh et al., 2007; Yerushalmi et al., 2006). The distance traveled by the bee in pixels is calculated by comparing the location on successive images on the camera field of view. Each camera recorded activity in 30 arenas (i.e., monitoring cages) on a single tray. Four cages, one on each tray, were left empty as controls, recording background noise.

### Analyses of circadian rhythms and sleep

We used the ClockLab circadian analysis software package (version 6; Actimetrics, Wilmette, IL) for all circadian rhythms and sleep analyses. We used 10-min bins to generate actograms and for the periodogram analyses. The χ2 periodograms were applied to the activity data collected on days 4–7 (Trial 1), days 4-9 (Trial 2), and days 4-10 (Trial 3) starting at 06:00 of the morning following transfer to the monitoring chamber. We used the ‘*Power*,’ calculated as the height of the periodogram peak relative to a significance threshold equal to *p* = 0.01, as an index for the strength of circadian rhythms (for more details, see Yerushalmi et al., 2006). Bees with a periodogram peak below the threshold line were assigned a zero-power value. The free-running period (*tau*) was determined as the period length below the prominent peak of the χ2 periodogram. The age of first showing circadian rhythmicity was determined as the first 3-day sliding window in which the periodogram analysis produced a statistically significant rhythm with a period between 20–28 h (Eban-Rothschild et al., 2011; Yerushalmi et al., 2006). The overall level of locomotor activity was determined using ClockLab analysis software and represented as counts. A sleep state was defined as a bout of 5 min or longer with no movement. This sleep proxy is based on detailed video analyses of the sleep-like behavior of individually isolated *B. terrestris* workers (Nagari et al., 2019). This index is similar to the sleep indices used for honey bees (Eban-Rothschild and Bloch, 2008;) and *Drosophila melanogaster* fruit flies (Shaw et al., 2000). The analyses of sleep were done using the ClockLab software, following preliminary analyses showing that the measurements are comparable to those obtained with the BeeSleep and Sleepograms algorithms that we used in previous studies (Eban-Rothschild and Bloch, 2008; Nagari et al., 2019)

### Assessing ovarian state

We thawed the stored bee samples and immersed them in bee saline (Huang et al., 1991) on a wax-filled dissecting plate under a stereomicroscope (Nikon SMZ645). We cut three incisions through the lateral and proximal-ventral abdominal cuticle using fine scissors and immersed the internal organs in honey bee saline. We used fine forceps to gently remove the ovaries into a drop of saline on a microscope slide and use the ocular ruler to measure the length of the four largest terminal oocytes form both ovarioles.

### Statistical Analyses

To assess the affect of JH manipulation on the Free Running Period (FRP, *Tau*), power, locomotor activity, the proportion of sleep, and oocyte size, we first tested whether each variable fits normal distribution, using the single variable Kolmogorov-Smirnov test. Based on the obtained results, we then determined whether the distribution of the abovementioned variables is the same across all three trials using either a one-way ANOVA for the normally or the Kruskal-Wallis test for the non-normally distributed variables. Given that these analyses revealed differences between the trials, we analyzed each trial separately and then combined the *p*-values of each trial to obtain an overall *p*-value. The combined *p*-values were calculated either by the Winer method of adding *t*’s (for parametric analyses) or by the Mosteller-Bush method of adding weighted *z*’s (for non-parametric analyses; see Rosenthal 1978). We used Cohen’s *d* as an approximation for the overall effect size in the pooled data. We next tested for each variable, whether the Control and Sham treatments are statistically different. We met this goal using either the parametric independent samples *t*-test or the non-parametric Mann-Whitney *U* test for each trial and then combine the results of each trial to obtain an overall *p*-value.

Given that for all variables tested, these two groups did not differ statistically, we used in subsequent analyses the Sham treatment group as the single control for the P-I and P-I+JH treatments. For assessing the treatment effect, we used pairwise comparisons between the Sham, P-I, and P-I+JH treatments, using either the parametric independent samples *t*-test or the non-parametric Mann-Whitney *U* test for each trial, as appropriate. We then combined the *p*-values of the three trials using either of the procedures mentioned above, and the final significance was determined by applying the Bonferroni correction for multiple comparisons.

For analyzing the rate of rhythm development, we considered only Trials 2 and 3; during days 4 to 8 (Trial 1, had only two 3-day sliding windows). In these two trials, each bee has five dependent observations, which are the Power value of the sliding windows on days 4 to 8. Thus, each bee can be represented by the slope of the linear regression of Power over time. Note that this step of the analysis does not require any assumptions of normality (since we do not test the significance of each slope). Next, we calculated the group means of the slopes of the individual bees for each Treatment. We further tested for normality of these slope samples. If these slope samples are normally distributed, we can obtain the combined *p*-value over both trials by using the Winer method of adding *t*’s. Given that *B. terrestris* workers develop circadian rhythms in locomotor activity with age (Yerushalmi et al., 2006), we expected the slopes to be positive, allowing us to present one-tailed *p*-values.

## Results

### The effect of JH on ovarian state

Given that previous studies show that JH is the major gonadotropin regulating oogenesis in *B. terestris* (Amsalem et al., 2014; Pandey et al., 2020; Shpigler et al., 2014; Shpigler et al. 2016), we used ovarian activity as a proxy for JH levels. The distribution of oocyte length did not differ from normal (Kolmogorov-Smirnov test, *p* = 0.134), but equality of variances among the trials was rejected (Levin’s test, *p* < 0.001). For the sake of consistency with the other analyses (see below), we treated here each trial separately, and then combined the results of each trial to obtain an overall *p*-value.

The Control and the Sham treatments did not differ statistically (independent samples *t*-test: Trial 1, *n*_Control_ = 19, *n*_Sham_ = 18, *t*_35_ = −0.837, *p* = 0.408; Trial 2, *n*_Control_ = 19, *n*_Sham_ = 20, *t*_37_ = 1.435, *p* = 0.160; Trial 3, *n*_Control_ = 19, *n*_Sham_ = 22, *t*_39_ = −0.228, *p* = 0.821). The combined significance level using the Winer method of adding *t*’s, was *z* = 0.208, with an overall *p* = 0.835, and an overall effect size = 0.007. Based on these analyses, we used only the Sham treatment as the control group, and carried out pairwise comparisons between Sham, P-I and P-I+JH treatments, using independent samples *t*-tests. The P-I treated bees have less developed ovaries compared to both the Sham and Replacement Therapy treatments in all three trials (**Fig. 2**.; independent samples *t*-test for comparing the means; *Sham vs. P-I:* Trial 1, *n*_Sham_ = 18, *n*_P-I_ = 16, *t*_23_ = 4.865, *p* < 0.001; Trial 2, *n*_Sham_ = 20, *n*_P-I_ = 22, *t*_28_ = 4.550, *p* < 0.001; Trial 3, *n*_Sham_ = 22, *n*_P-I_ = 7, *t*_27_ = 2.527, *p* = 0.018; overall *z* = 6.622, *p* < 0.001, effect size = 0.850; *P-I vs. P-I+JH:* Trial 1, *n*_P-I_ = 16, *n*_P-I+JH_ = 24, *t*_37_ = −7.632, *p* < 0.001; Trial 2, *n*_P-I_ = 22, *n*_P-I+JH_ = 20, *t*_32_ = −8.373, *p* < 0.001; Trial 3, *n*_P-I_ = 7, *n*_P-I+JH_ = 15, *t*_20_ = −4.397, *p* < 0.001; overall *z* = −11.345, *p* < 0.001, effect size = 1.704 (*p*-values in bold are significant after Bonferroni correction). These analyses of ovarian state confirm that our treatments effectively manipulated JH levels. There were no consistent differences between the Sham vs. P-I+JH treatments indicating that our replacement therapy successfully reverted the effect of JH reduction by the P-I treatment (Trial 1, *n*_Sham_ = 18, *n*_P-I+JH_ = 24, *t*_40_ = −0.674, *p* = 0.504; Trial 2, *n*_Sham_ = 20, *n*_P-I+JH_ = 20, *t*_38_ = −2.042, *p* = 0.048; Trial 3, *n*_Sham_ = 22, *n*_P-I+JH_ = 15, *t*_33_ = −1.321, *p* =0.198; overall *z* = −2.267, *p* = 0.023, effect size = 0.266).

**Figure 2.**
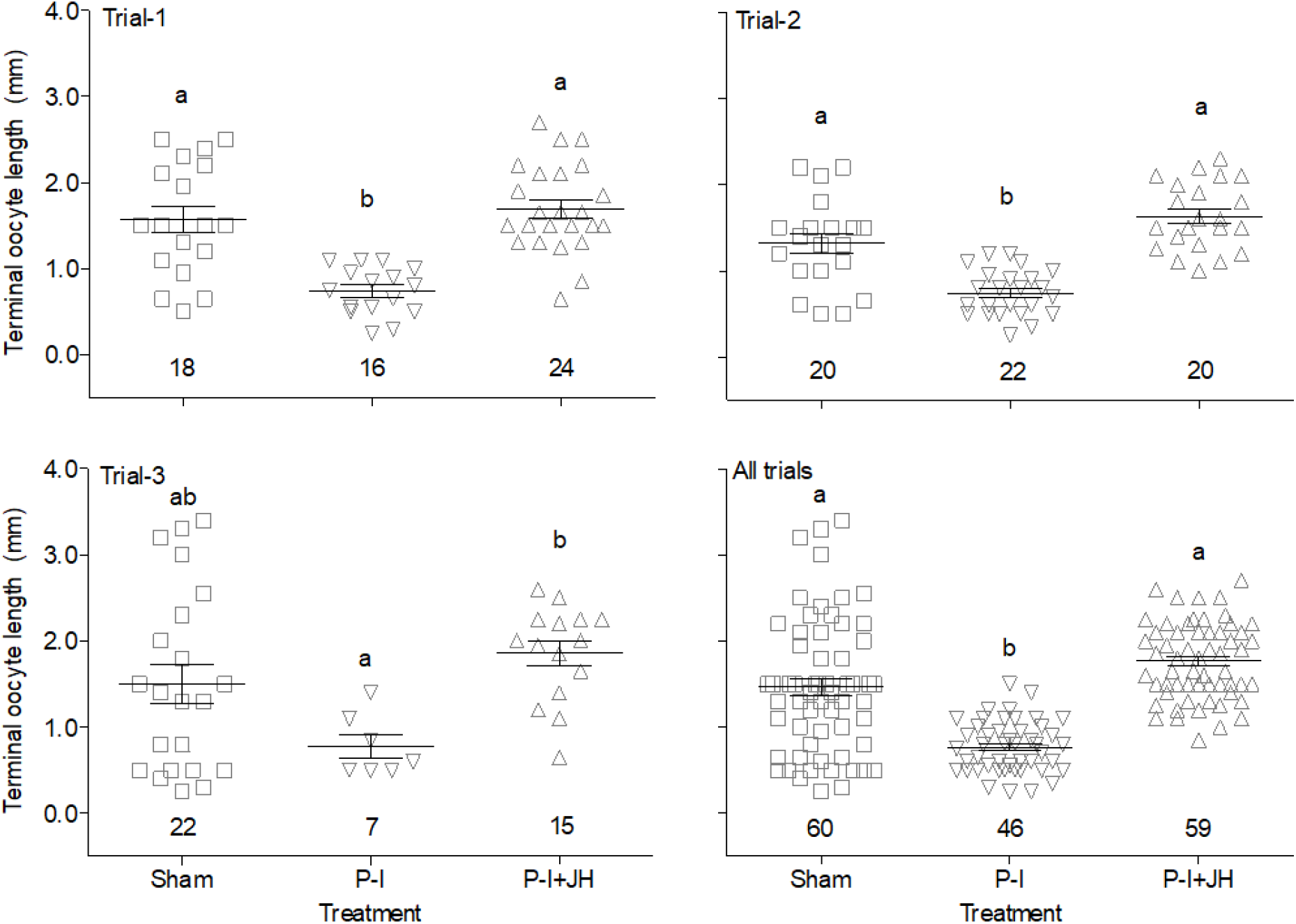
The effect of manipulating JH levels on ovarian state. Sham - Sham treatment with only the vehicles used to deliver the drugs; P-I – Precocene-I treatment to reduce JH levels; P-I+JH - replacement therapy with JH-III, the natural JH of bumble bees. Open symbols depict values for individual bees, the horizontal lines and error bars show the mean ± SE, and the sample size is reported just above the x-axis line. Treatments marked with different letters are statistically different using independent samples t-test followed by Bonferroni correction. For the combined analysis (“All trials” bottom right panel), we used the Winer method of adding *t’s* after applying the Bonferroni correction.

### The effect of JH on the overall level of activity

For each bee, we calculated the mean activity per hour during days 1 to 4. The distribution of these individual means did not differ from normal distribution (Kolmogorov-Smirnov test for normality: *p* = 0.239, *p* = 0.334 and *p* = 0.198 for Trials 1, 2 and 3, respectively). The Control and the Sham treatments did not differ statistically (independent samples *t*-test: Trial 1, *n*_Control_ = 19, *n*_Sham_ = 19, *t*_30_ = −0.513, *p* = 0.611; Trial 2, *n*_Control_ = 19, *n*_Sham_ = 20, *t*_37_ = 1.681, *p* = 0.101; Trial 3, *n*_Control_ = 19, *n*_Sham_ = 22, *t*_31_ = 0.012, *p* = 0.991). Using the Winer method of adding *t*’s, the combined *z* = 0.660, with an overall *p* = 0.509, and an overall effect size = 0.057. Thus, we used only the Sham treatment as the control group, and carried out pairwise comparisons between Sham, P-I and P-I+JH treatments, using independent samples *t*-tests. tests. *Sham vs. P-I:* Trial 1, *n*_Sham_ = 19, *n*_P-I_ = 16, *t*_33_ = 0.658, *p* = 0.515; Trial 2, *n*_Sham_ = 20, *n*_P-I_ = 22, *t*_40_ = - 0.411, *p* = 0.683; Trial 3, *n*_Sham_ = 22, *n*_P-I_ = 7, *t*_27_ = −0.314, *p* = 0.756; overall *z* = −0.037, *p* = 0.970, effect size = 0.067. *Sham vs. P-I+JH:* Trial 1, *n*_Sham_ = 19, *n*_P-I+JH_ = 24, *t*_41_ = −1.293, *p* = 0.203; Trial 2, *n*_Sham_ = 20, *n*_P-I+JH_ = 20, *t*_38_ = 0.200, *p* = 0.842; Trial 3, *n*_Sham_ = 22, *n*_P-I+JH_ = 15, *t*_35_ = 1.398, *p* = 0.171; overall *z* = 0.171, *p* = 0.864, effect size = 0.021. *P-I vs. P-I+JH:* Trial 1, *n*_P-I_ = 16, *n*_P-I+JH_ = 24, *t*_38_ = −1.820, *p* = 0.077; Trial 2, *n*_P-I_ = 22, *n*_P-I+JH_ = 20, *t*_40_ = 0.562, *p* = 0.577; Trial 3, *n*_P-I_ = 7, *n*_P-I+JH_ = 15, *t*_20_ = 1.118, *p* = 0.277; overall *z* = −0.078, *p* = 0.938, effect size = 0.080. **Table 2** summarizes the means of the individual mean activity for each of the three trials.

**Table 2.** Manipulating JH titers did not affect the overall level of locomotor activity. The values are mean ± se, with sample size in parentheses.

| Treatment | Trial 1 | Trial 2 | Trial 3 |
| --- | --- | --- | --- |
| Sham | 704 ± 76 (19) | 512 ± 48 (20) | 704 ± 77 (22) |
| P-I | 633 ± 77 (16) | 544 ± 61 (22) | 761 ± 213 (7) |
| P-I+JH | 858 ± 87 (24) | 497 ± 57 (20) | 529 ± 102 (15) |

### The influence of JH on the strength of circadian rhythms

We used the Power as an index for the strength of circadian rhythms in locomotor activity. Given that the Power was not normally distributed (Kolmogorov-Smirnov test, *p* < 0.001), we used non-paramteric statistics. The distribution of Power differed among the three trials (Kruskal-Wallis test, *p* < 0.001). Therefore, we treated each trial separately, and then combined the *p*-value of each trial to obtain a combined *p*-value. We used the Sham as our control group following a preliminary test showing no difference between the Control and the Sham treatments in all three trials (Mann-Whitney *U* test: Trial 1, *n*_Control_ = 19, *n*_Sham_ = 19, *z* = −0.394, *p* = 0.707; Trial 2, *n*_Control_ = 19, *n*_Sham_ = 20, *z* = −0.141, *p* = 0.895; Trial 3, *n*_Control_ = 19, *n*_Sham_ = 22, *z* = −0.013, *p* = 0.995; Mosteller-Bush method of adding weighted *z*’s, *z* = −0.131, an all trials combined, significance level *p* = 0.896, and an effect size = 0.011). In all three trials there was a consistent trend of attenuated circadian rhythms for the P-I treated bees compared to the other groups, but the individual trials the differences were statistically significant after Bonferroni only in the first trial for the comparison of the P-I and Sham treatments. Nevertheless, the differences were statistically significant after correction in a pooled analyses that combined the p-values of all three trials (**Fig. 3**; *Sham vs. P-I:* Trial 1, *n*_Sham_ = 19, *n*_P-I_ = 16, *z* = 2.563 *p* = 0.009; Trial 2, *n*_Sham_ = 20, *n*_P-I_ = 22, *z* = 2.242, *p* = 0.025; Trial 3, *n*_Sham_ = 22, *n*_P-I_ = 7, *z* = 0.561, *p* = 0.600; All trials combined, *z* = 3.234, *p* = 0.001, effect size = 0.554; *Sham vs. P-I+JH:* Trial 1, *n*_Sham_ = 19, *n*_P-I+JH_ = 24, *z* = 1.095, *p* = 0.279; Trial 2, *n*_Sham_ = 20, *n*_P-I+JH_ = 20, *z* = −0.839, *p* = 0.409; Trial 3, *n*_Sham_ = 22, *n*_P-I+JH_ = 15, *z* = 0.449, *p* = 0.663 (Trial 3); All trials combined, *z* = 0.434, *p* = 0.664, effect size = 0.006; *P-I vs. P-I+JH:* Trial 1, *n*_P-I_ = 16, *n*_P-I_+JH = 24, *z* = −1.453, *p* = 0.149; Trial 2, *n*_P-I_ = 22, *n*_P-I+JH_ = 20, *z* = −2.040, *p* = 0.042; Trial 3, *n*_P-I_ = 7, *n*_P-I+JH_ = 15, *z* = −0.390, *p* = 0.729; all trials combined, *z* = −2.456, *p* = 0.014, effect size = 0.459).

**Figure 3.**
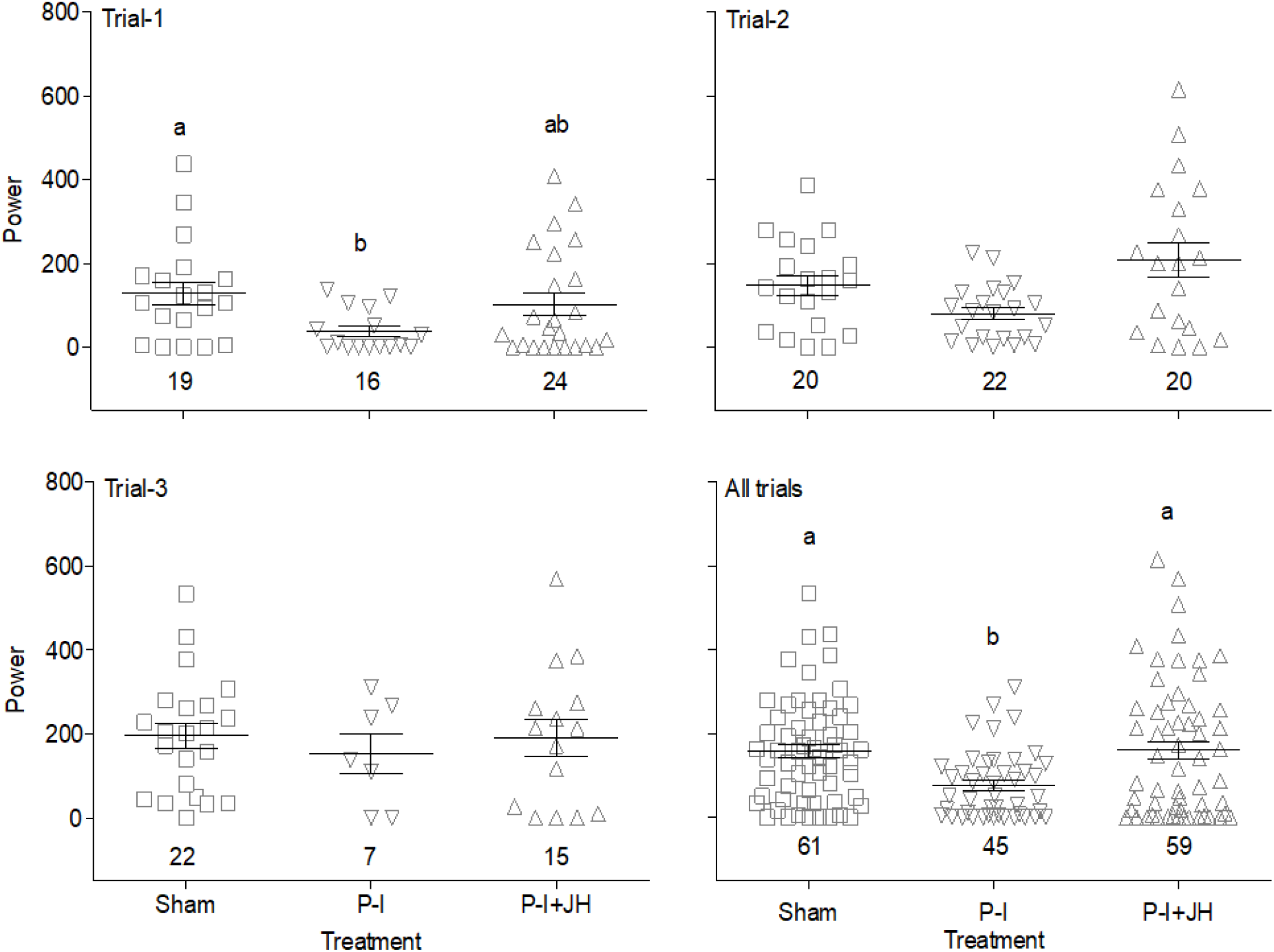
The effect of manipulating JH levels on the strength of circadian rhythm in locomotor activity. The Y-axis shows the Power, which we used as an index for the strength of circadian rhythm (arbitrary units). Treatments marked with different letters are statistically different after applying the Bonferroni correction in independent samples t-test for each trial, and in Mosteller-Bush method, in the pooled analyses (‘All trials’, bottom right) panel. See Figure 2 for additional details.

### The influence of JH on the free-running period (*Tau*)

Given that the distribution of *Tau* was not normal (Kolmogorov-Smirnov test, *p* = 0.001), we used non-parametric statistics. The distribution differs for the three trials (Kruskal-Wallis test, *p* < 0.001), and therefore we treated each trial separately, and then combined the obtained *p*-values to an overall *p*-value. We used the Sham as a single control group following a preliminary analysis showing that the FRP is not different between the Control and the Sham treatments in all three trials (Mann-Whitney *U* test for each trial: Trial 1, *n*_Control_ = 19, *n*_Sham_ = 16, *z* = −0.695, *p* = 0.497; Trial 2, *n*_Control_ = 16, *n*_Sham_ = 17, *z* = 0.525, *p* = 0.611; Trial 3, *n*_Control_ = 18, *n*_Sham_ = 21, *z* = 1.089, *p* = 0.283) as well as in a Mosteller-Bush method for combining all trials combined (weighted *z*’s, we got *z* = 0.573, *p* = 0.567, effect size = 0.110). The *Tau* did not differ between the Sham, P-I, and Replacement Therapy treatment groups in any of the three trials or in the pooled analysis (**Fig. 4**.; Mann-Whitney *U* test for pairwise comparisons: Trial 1, *Sham vs. P-I: n*_Sham_ = 16, *n*_P-I_ = 8, *z* = 0.858, *p* = 0.408; Trial 2, *n*_Sham_ = 17, *n*_P-I_ = 19, *z* = 0.666, *p* = 0.515; Trial 3, *n*_Sham_ = 21, *n*_P-I_ = 5, *z* = −1.374, *p* = 0.180; All trials combined, *z* = 0.175, *p* = 0.861, effect size = 0.006. Sham vs. P-I+JH: Trial 1, *n*_Sham_ = 16, *n*_P-I+JH_ = 17, *z* = 1.459, *p* = 0.149; Trial 2, *n*_Sham_ = 17, *n*_P-I+JH_ = 18, *z* = −0.976, *p* = 0.338; Trial 3, *n*_Sham_ = 21, *n*_P-I+JH_ = 12, *z* = −2.196, *p* = 0.027; All trials combined, *z* = −1.002, *p* = 0.316, effect size = 0.135. *P-I vs. P-I+JH:* Trial 1, *n*_P-I_ = 8, *n*_P-I+JH_ = 17, *z* = −0.321, *p* = 0.765; Trial 2, *n*_P-I_ = 19, *n*_P-I+JH_ = 18, *z* = −1.933, *p* = 0.054; Trial 3, *n*_P-I_ = 5, *n*_P-I+JH_ = 12, *z* = −0.159, *p* = 0.898; All trials combined, *z* = −1.721, *p* = 0.085, effect size = 0.115).

**Figure 4.**
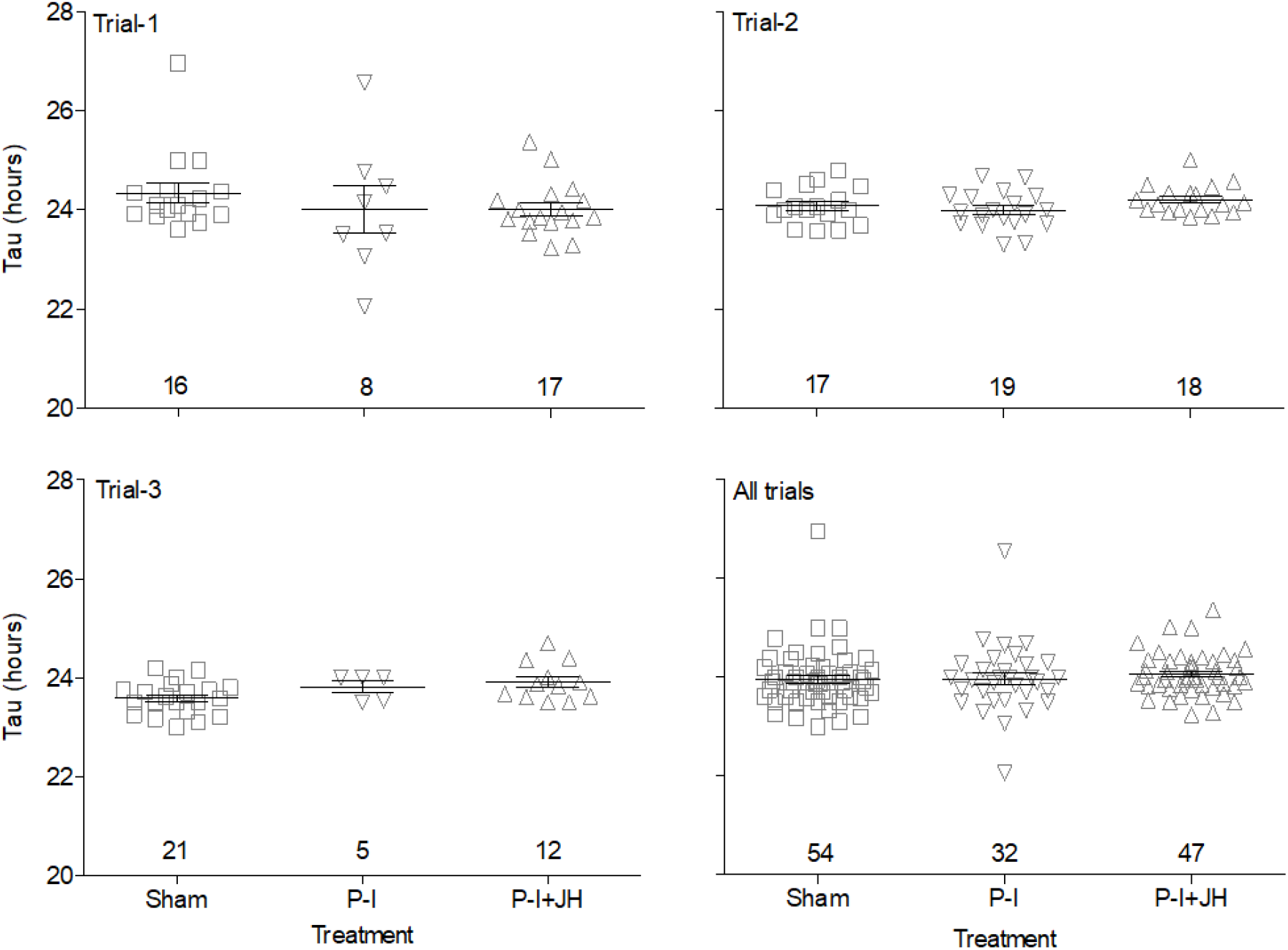
The affect of manipulating JH levels on the free running period (*tau*). The Y-axis shows the circadian period length under constant conditions. See Figure 2 for additional details.

### The influence of JH on the ontogeny of circadian rhythms

For this experiment we considered only the second and third trials during days 4 to 8; Trial 1 was not included because we monitored bees for only seven days, which is not sufficient for developmental analyses. For each bee, we measured the Power for consecutive 3-day sliding window, and used these values to calculate the slope of the linear regression of Power as a function of age (days 4 to 9). The slope values did not differ from normal distribution (Kolmogorov-Smirnov test for normality: *p* = 0.241 and *p* = 0.591 for Trials 2 and 3, respectively). **Table 3** summarizes the means of these slopes for each trial followed by the combined one-tailed *p*-value and the effect size for each treatment. The combined *p*-value was obtained by applying the Winer method of adding *t*’s. Whereas all sample means are positive, only the combined *p*-value for the replacement therapy (P-I+JH) treatment is statistically significant.

**Table 3.**
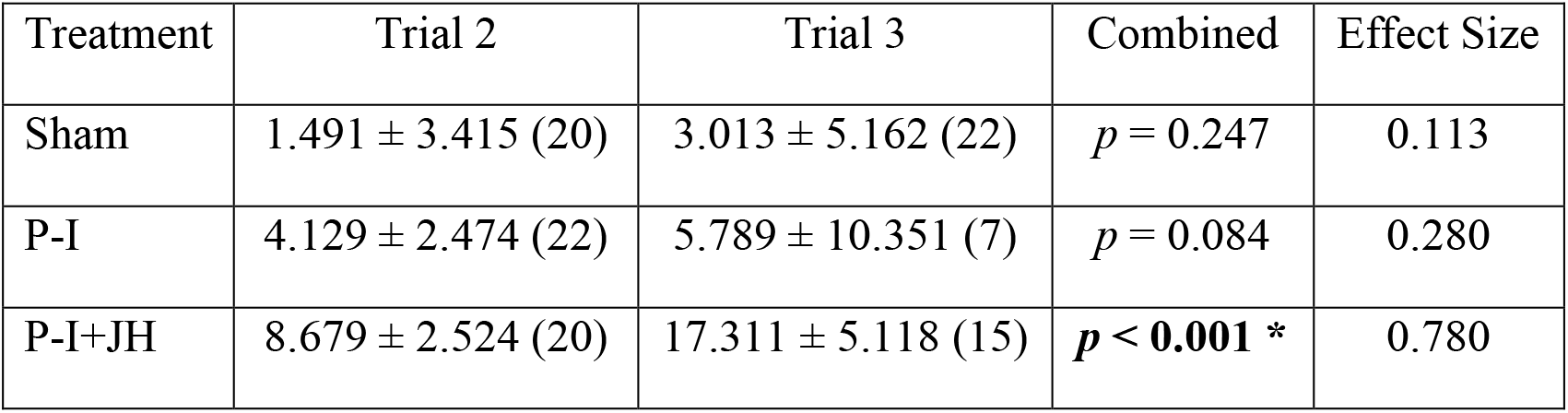
The slopes of the regression lines for the strength of circadian rhythms (Power) as a function of age. The values are means ± se, the sample size in parentheses. The fourth column presents the combined *p*-value obtained by applying the Winer method. The rightmost column presents the combined effect size for each treatment. An asterisk denotes statistically significance difference in a one-tailed test.

We next used *t*-tests to test the influence of JH manipulation on the slope of an increase in the strength of circadian rhythms as a function of age. These analyses are summarized in **Table 4**, showing significantly faster rhythms development for replacement therapy treated bees (P-I+JH) compared to the Sham control. There was a similar trend of faster development for the Replacement Therapy compared to P-I treatment, but the *p*-value (*p*=0.051) did not cross our statistical significance threshold after Bonferroni correction (*p*<0.017). These analyses provide weak support for the hypothesis that JH influences the development of circadian rhythms in young bumble bees.

**Table 4.** The influence of JH manipulation on the rate of development of circadian rhythms in locomotor activity. The table shows the *t* statistic of the comparisons between the mean slopes for each pair of treatments. A negative sign indicates that the mean of the first treatment is smaller than that of the second treatment in comparison. The fourth and rightmost columns summarized the combined *p*-value, and effect size, respectively. An asterisk denotes statistically significance difference in a one-tailed test.

| Comparison | Trial 2 | Trial 3 | Combined | Effect Size |
| --- | --- | --- | --- | --- |
| Sham vs. P-I | $t_{40} = -0.634$ | $t_{27} = -0.256$ | $p = 0.271$ | 0.087 |
| Sham vs. P-I+JH | $t_{38} = -1.692$ | $t_{35} = -1.893$ | $p = \mathbf{0.007}^*$ | 0.393 |
| P-I vs. P-I+JH | $t_{40} = -1.285$ | $t_{20} = -1.126$ | $p = 0.051$ | 0.346 |

### The influence of JH on the daily sleep amount of individually isolated bees

For each bee, we calculated the mean proportion of time asleep (inactivity bouts of 5 min or more) during days 1 to 4. The distribution of these individual means did not differ from normal distribution (Kolmogorov-Smirnov test for normality: *p* = 0.812, *p* = 0.450 and *p* = 0.256 for Trials 1, 2 and 3, respectively). The Control and the Sham treatments did not differ statistically in any of the trials (independent samples *t*-test: Trial 1, *n*_Control_ = 19, *n*_Sham_ = 19, *t*_36_ = 0.329, *p* = 0.744; Trial 2, *n*_Control_ = 19, *n*_Sham_ = 20, *t*_37_ = −0.628, *p* = 0.534; Trial 3, *n*_Control_ = 19, *n*_Sham_ = 22, *t*_39_ = −0.505, *p* = 0.616), as well as in a pooled analysis using the Winer method of adding *t*’s (combined *z* = −0.452, overall *p* = 0.652, overall effect size = 0.053). Thus, we used only the Sham treatment as the control group, and carried out pairwise comparisons between Sham, P-I and P-I+JH treatments, using independent samples *t*-tests. *Sham vs. P-I*: Trial 1, *n*_Sham_ = 19, *n*_P-I_ = 16, *t*_33_ = −1.530, *p* = 0.136; Trial 2, *n*_Sham_ = 20, *n*_P-I_ = 22, *t*_40_ = 1.041, *p* = 0.304; Trial 3, *n*_Sham_ = 22, *n*_P-I_ = 7, *t*_27_ = −0.565, *p* = 0.577; overall *z* = −0.589, *p* = 0.556, effect size = 0.113. *Sham vs. P-I+JH:* Trial 1, *n*_Sham_ = 19, *n*_P-I+JH_ = 24, *t*_41_ = −0.382, *p* = 0.704; Trial 2, *n*_Sham_ = 20, *n*_P-I+JH_ = 20, *t*_31_ = −1.443, *p* = 0.159; Trial 3, *n*_Sham_ = 22, *n*_P-I+JH_ = 15, *t*_35_ = −0.213, *p* = 0.832; overall *z* = −1.112, *p* = 0.266, effect size = 0.186. *P-I vs. P-I+JH:* Trial 1, *n*_P-I_ = 16, *n*_P-I+JH_ = 24, *t*_38_ = 1.095, *p* = 0.280; Trial 2, *n*_P-I_ = 22, *n*_P-I+JH_ = 20, *t*_31_ = −2.474, *p* = 0.019; Trial 3, *n*_P-I_ = 7, *n*_P-I+JH_ = 15, *t*_20_ = 0.354, *p* = 0.727; overall *z* = −0.570, *p* = 0.569, effect size = 0.053. The amount of sleep for the Sham, P-I and replacement therapy treated bees are summarized in **Table 5**.

**Table 5.** The influence of JH on the amount of sleep of individually isolated workers. The table shows means ± se, with the sample size in parentheses. None of the comparison is statistically significant after Bonferroni correction (see text for details).

| Treatment | Trial 1 | Trial 2 | Trial 3 |
| --- | --- | --- | --- |
| Sham | 0.366 ± 0.024 (19) | 0.401 ± 0.031 (20) | 0.291 ± 0.026 (22) |
| P-I | 0.429 ± 0.034 (16) | 0.350 ± 0.037 (22) | 0.322 ± 0.056 (7) |
| P-I+JH | 0.381 ± 0.028 (24) | 0.453 ± 0.019 (20) | 0.300 ± 0.034 (15) |

## Discussion

JH is the major gonadotropin in many insects, including bumble bees, in which it regulates physiological processes such as oogenesis, vitellogenesis, oogenesis, and wax production, and behaviors such as dominance and aggression. Little is known, on the influences of JH on circadian rhythms and sleep. We used a pharmacological approach to test the hypothesis that JH influences circadian rhythms or sleep in individually isolated bumble bee workers. Our measurements of ovarian activity confirm that our manipulations were effective; bees treated with P-I to reduced JH levels had inactive ovaries, and this effect was fully reverted by replacement therapy with JH-III, the natural JH of this species (Bloch et al., 2000, 1996). We found that bees with reduced JH levels have weaker circadian rhythms compared to control bees, an effect that was reverted by replacement therapy. The slope of circadian rhythm strengthening with age was more moderate for bees with reduced JH titers suggesting that the developmental effects of JH may contribute to its influence on the strength of circadian rhythms. To our knowledge, our results provide the strongest support for the hypothesis that JH augments circadian rhythms in an insect.

There is good evidence that gonadotropic hormones influence circadian rhythms in vertebrates (reviewed in Bedrosian et al., 2016; Gamble et al., 2014), which makes functional sense given that gonadotropins coordinate processes in many tissues involved in reproduction. The evidence for modulation of circadian processes by gonadotropic hormones in insects is more ambiguous (see Introduction; Bloch et al., 2013). Here we show that JH, the major gonadotropic hormone in the bumble bee *B. terrestris* (Pandey et al., 2020; Shpigler et al., 2014, 2016), influences the development and strength of circadian rhythms. Notably, these findings contrast with a study with honey bees in which allatectomy and replacement therapy manipulations did not affect circadian rhythms in locomotor activity in individually isolated workers (Bloch et al., 2002). This incongruity is consistent with other evidence that JH does not function as a gonadotropin in honey bees and overall, has different functions in these two social bees (Bloch, G., Wheeler, D., Robinson, 2002; Hartfelder, 2000; Robinson and Vargo, 1997).

Our circadian analyses suggest that the most significant influence of JH manipulation was on the strength of circadian rhythms (**Fig. 3**), with an overall no effect on the endogenous circadian period (**Fig. 4**). Hormones may affect the strength of circadian rhythms in locomotor activity by several not mutually exclusive mechanisms. First, JH may act downstream of the circadian clock to regulate locomotor activity such that oscillations in circulating JH levels control daily changes in the level of locomotor activity. There is indeed evidence for circadian modulation of JH in insects, including honey bees (reviewed in Bloch et al., 2013). Moreover, in some insects, including honey bees, JH augments metabolic rates (Denlinger et al., 1984; Garcera et al., 1991; Sláma and Ilona, 1979; Sullivan et al., 2003). In some species such as the cockroach *B. germanica* there is even direct evidence that JH augments the overall level of activity (Lin and Lee, 1998). However, we did not find a reduction in overall locomotor activity in P-I treated bees or a significant increase in the replacement therapy (**Table 2**). JH similarly did not affect the level of locomotor activity in honey bees (Bloch et al., 2002). The main inconsistency of this model with our finding is that it implies that insects with no JH will not show circadian rhythms. By contrast, we found that bees treated with P-I, that is assumed to disable the CA, nevertheless, show robust rhythms (although attenuated relative to control bees). Moreover, a single acute, not cyclic treatment with JH, fully recovered the rhythm attenuating effect of P-I. A second possible explanation for our findings is that oscillations in JH titers augment, but are not imperative for circadian rhythms in locomotor activity. According to this model, central clocks in the brain concurrently influence both locomotor activities controlling centers and JH signaling (e.g., regulating JH biosynthesis in the CA; Bloch et al., 2013). Circadian rhythms in JH signaling in turn act on locomotor activity centers to augment exciting circadian rhythms in locomotor activity. As detailed above for the first hypothesis, it is not easy to reconcile this presumed mechanism with our finding that a single acute treatment with JH at early age successfully reverted the rhythm attenuating affect of the P-I treatment. A third hypothesis states that JH has activational effects regulating functions in the central circadian network or downstream output pathways controlling locomotion. This idea is consistent with studies with rodents showing that sex steroids affect various circadian parameters of locomotor activity, including the phase, FRP, amplitude (i.e., strength), and splitting of locomotor activity rhythms (Daan et al., 1975; Morin, 1980; Morin et al., 1977; reviewed in Hatcher et al., 2020). Many of these effects could be attained by acute pharmacological treatments acting on hormone receptors indicating that the hormonal influence is activational rather than developmental (e.g., Karatsoreos et al., 2011; Model et al., 2015). In addition, treatments with steroid hormones were shown to regulate the expression of genes that are important for the generation or expression of circadian rhythms within specific cells, or by enhancing coupling within the timekeeping system (e.g., Karatsoreos et al., 2011; Nakamura et al., 2008, 2005; reviewed in Hatcher et al., 2020). Activational effects may also be mediated by the hormones acting on peripheral clocks or effector tissues (He et al., 2007; Sellix et al., 2004). The last hypothesis is that JH regulates the organization (development) of the circadian system. In mammals, it is well-established that sex steroid hormones have various organizational effects regulating the development of the circadian neural network and the expression of estrogen and androgen receptors which underlie sexual dimorphism in circadian activity (Hagenauer et al., 2011a, 2011b; Hummer et al., 2012; Melo et al., 2010; Royston et al., 2016; Sellix et al., 2013; for a recent review see Hatcher et al., 2020). Our findings that a single acute manipulation of JH levels shortly after adult eclosion from the pupa, resulted in relatively long term effects on the power of circadian rhythms best fit with the forth hypothesis stating that the influence of JH on circadian rhythms is mostly organizational. The steeper age-related increase in the strength of circadian rhythms in bees subjected to replacement therapy provides additional support for this hypothesis (Tables 3 and 4). However, it should be noted that activational effects (third hypothesis above) may also account for at least some of the effects of JH on circadian rhythms.

Our findings that JH augments circadian rhythms in workers are puzzling because reproduction in *B. terrestris*, that is regulated by JH, is typically associated with activity around the clock and attenuated circadian rhythms in locomotor activity. Bumble bees gynes show robust circadian rhythms but later switch to activity around the clock when they establish colonies and reproduce (Eban-Rothschild et al., 2011). Similarly, nest workers, that typically have better-developed ovaries and higher JH titers (Shpigler et al., 2016; van Honk et al., 1981) are typically active around the clock in the presence of brood (Nagari et al., 2019; Yerushalmi et al., 2006). A plausible explanation for this apparent discrepancy is that we studied individually isolated bees, and rhythm manifestation in nurse bees is context dependent (Eban-Rothschild and Bloch, 2012b; Yerushalmi et al., 2006). Our finding that JH augments circadian rhythms in individually isolated workers does seem to reflect organizational effects on the development or integration of the circadian systems. The faster development of the circadian system in bees with high JH titers are not manifested in the colony because nurse workers and egg-laying queens are typically active around the clock in the presence of brood (Fig. 1.2 in Eban-Rothschild and Bloch, 2012b). Perhaps, a functional circadian clock system is regulated by a gonadotropin because it improves the coordination of internal processes related to reproduction. This idea is only a speculation at this stage. However, it is consistent with studies with mammals in which circadian clocks in the gonads and endocrine organs are imperative for fertility, and are sensitive to gonadotrophic hormones (reviewed in Sellix and Menaker, 2011). This may also be the case in male flies: *Drosophila melanogaster* flies with mutations in each of the clock genes *Period, Timeless, Cycle*, and *Clock*, show reduced fertility. The low-fertility phenotype was reversed in flies with rescued *Period* or *Timeless* function confirming the importance of these canonical clock genes. Further crosses between mutant and wild-type flies indicate that clock mutations compromised the fertility of both the male and female flies (Beaver et al., 2002). In males, peripheral circadian oscillators in the testis vas deferens complex are necessary for fertility (Beaver et al., 2002). However, in females, in which clock gene levels do not cycle in the ovary, the effects of clock gene mutations on fertility is due to non-circadian functions of the *Period* and *Timeless* in the ovary (Beaver et al., 2003). There is also evidence that JH interacts with clock genes in other insects. For example, in the linden bug, *Pyrrhocoris apterus*, JH regulates gene expression in the gut through interactions of its receptor Met with the circadian proteins Clock and Cycle (Bajgar et al., 2013).

Additional studies with bees with mutated or downregulated clock genes are necessary for testing if JH regulation of circadian clocks is involved in the reproductive physiology of bumble bees, or that the effects that we report here are related to other processes that we did not explore.

## Acknowledgments

This study was supported by grants from the US-Israel Binational Agricultural Research and Development Fund (BARD, number IS-4418-11, and IS-5077-18, to G.B.) and The Planning and Budgeting Committee (PBC) Fellowship Program for Outstanding Chinese and Indian Post-Doctoral Fellows (to A.P).

